# An adaptive optics module for deep tissue multiphoton imaging *in vivo*

**DOI:** 10.1101/2020.11.25.397968

**Authors:** Cristina Rodríguez, Anderson Chen, José A. Rivera, Manuel A. Mohr, Yajie Liang, Wenzhi Sun, Daniel E. Milkie, Thomas G. Bifano, Xiaoke Chen, Na Ji

## Abstract

Understanding complex biological systems requires visualizing structures and processes deep within living organisms. We developed a compact adaptive optics module and incorporated it into two- and three-photon fluorescence microscopes, to measure and correct tissue-induced aberrations. We resolved synaptic structures in deep cortical and subcortical areas of the mouse brain, and demonstrated high-resolution imaging of neuronal structures and somatosensory-evoked calcium responses in the mouse spinal cord at unprecedented depths *in vivo*.

---

Imaging living organisms with subcellular resolution is key for understanding biological systems. Two-photon (2P) fluorescence microscopy is an essential tool for observing cells and biological processes under physiological conditions deep inside tissue. With longer excitation wavelength experiencing reduced tissue scattering, combined with the suppression of out-of-focus excitation due to the 3^rd^-order nonlinear optical process, three-photon (3P) fluorescence microscopy further extends the imaging depth in opaque tissues^1–3^. For both 2P and 3P microscopy beyond superficial depths, however, living biological tissues aberrate the wavefront of the excitation light, leading to an enlarged excitation focus of diminished focal intensity, ultimately limiting the *in vivo* imaging performance to cellular resolution at depth.

In the mouse central nervous system, reliably imaging the structure and function of neurons at subcellular resolution requires that the distortion of the excitation light, accumulated as it travels through tissues, be measured and corrected by adaptive optics (AO) methods^4–7^. How aberrations are measured is the main factor distinguishing the different implementations of AO in microscopy. While direct wavefront measurement methods have high measurement speed, they require separate wavefront sensors and correctors. In addition, the introduction of exogenous near-infrared emitting dyes is required to maintain their performance in highly scattering samples such as tissues^8,9^. Albeit slower, indirect wavefront sensing methods are well suited for opaque media. Using two wavefront modulation devices, an indirect wavefront sensing method based on frequency multiplexed aberration measurement successfully achieved 2P imaging of submicrometer-sized dendritic spines in layer 5 of the mouse primary visual cortex^10^. However, this method was limited to a narrow band of wavelengths and slow measurement times by the liquid-crystal spatial light modulator used for aberration correction. Here, we report a compact AO module for multiphoton microscopy, composed of a novel, high-speed segmented deformable mirror (DM), two lenses, and a field stop, that overcomes the aforementioned limitations. We demonstrate high-resolution 2P and 3P fluorescence imaging in a variety of optically challenging locations in the mouse central nervous system. With AO, we were able to resolve fine neuronal processes and synapses (e.g., dendritic spines) in deep cortical layers as well as subcortical areas of the mouse brain *in vivo* – features otherwise invisible without aberration correction. We further achieved high-resolution *in vivo* imaging of neuronal structures and somatosensory-evoked calcium responses in the mouse spinal cord at depths that to our knowledge have not been reported before.

The frequency multiplexed aberration measurement^10^ and the subsequent correction were implemented solely by the DM (Methods). We first determined the phase gradient values to be added to each DM segment such that the beamlets reflecting off them would converge to a single point on the focal plane. To achieve this, we formed a stationary reference focus using half of the beamlets and raster-scanned the other half of the beamlets around this focus by tip-tilting the corresponding DM segments. At each set of phase gradients (i.e., produced by tipping and tilting the mirror segments leading to beamlet displacements on the focal plane), we recorded the fluorescence signal variation while modulating the phase or intensity of the scanned beamlets at distinct ~kHz frequencies using the DM segments. The interference strength of each beamlet with the reference focus was obtained by Fourier transforming the signal trace and recording the Fourier magnitude at its modulation frequency. The phase gradients giving rise to maximal interference strength for these beamlets were then calculated. Swapping the scanned and the stationary DM segments and repeating the phase gradient measurement, we obtained the wavefront gradients required to converge all beamlets to the same focus. We then measured the phase of each beamlet directly following a similar modulation-based procedure^11^, again taking advantage of the high-speed DM segments. For larger aberrations, the whole procedure may be repeated for several iterations to achieve optimal aberration correction. The final corrective wavefront was then applied to the DM for aberration correction. After reflecting off the DM and traveling through the sample, all beamlets converged around a common spot with the same phase so that they constructively interfered to form a diffraction-limited focus. In contrast to previous work^10^, where a slow DM or digital micromirror device was used for frequency multiplexed modulation, and a spatial light modulator for aberration measurement and correction, using a single high-speed DM both simplifies our module and enables faster (~4×) measurement time, high power throughput, along with polarization- and wavelength-independent operation, which allowed us to use the same module for 2P and 3P fluorescence microscopy.

We incorporated our AO module into a homebuilt 2P fluorescence microscope by placing it between the excitation laser and the microscope (Supplementary Fig. 1a). We first validated its performance in correcting artificial aberrations using the signal from fluorescent features of different sizes. For one representative artificial aberration (Supplementary Fig. 2), both intensity and phase modulation successfully enabled corrections that substantially increased signal and improved resolution of fluorescent bead images, with higher signal recovery for smaller beads (23.7× for 0.5-μm-diameter bead versus 3.7× for 10-μm-diameter bead). We next applied the AO module to high-resolution *in vivo* imaging. In one example, we imaged GFP-labeled myotomes in the mid-trunk of a young zebrafish larva, using a 920 nm laser excitation wavelength (Fig. 1a-c). At an imaging depth of 110 μm below the surface, the aberration introduced by this sample, mostly astigmatism resulting from the larva’s highly curved cylindrical-like surface, led to images of low resolution and contrast (Fig. 1a). Aberration correction resulted in a ~2-fold increase of 2P fluorescence signal and most notably, a substantial (~3.6×) improvement in image contrast.

**Fig. 1.**
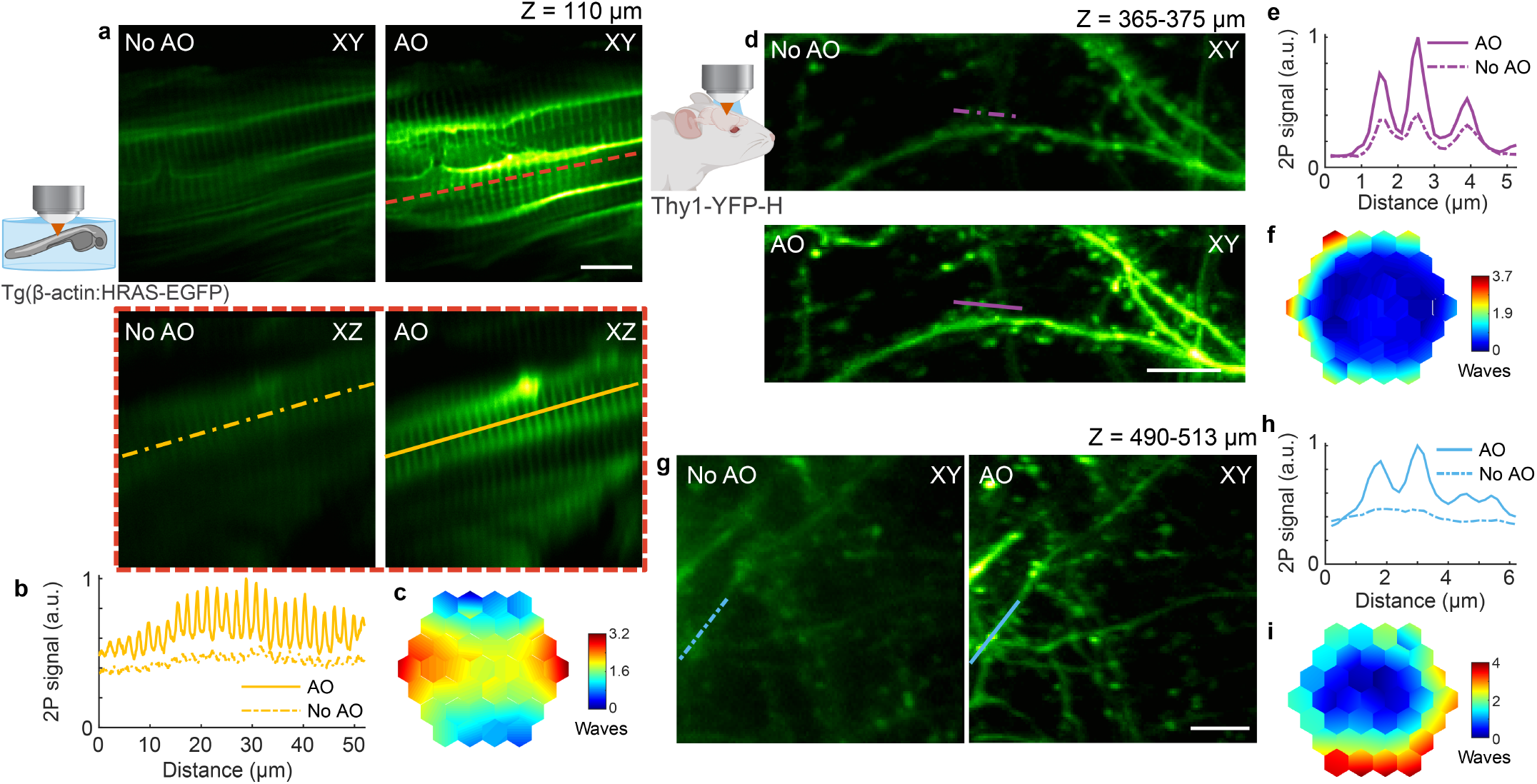
AO improves *in vivo* 2P imaging of myotomes in zebrafish larva and neuronal structures in the mouse brain. **a**, Lateral and axial (along red dashed line) images of myotomes in the mid-trunk of a 4-day-old zebrafish larva, at an imaging depth of 110 μm from the surface, without and with AO (phase modulation). Post-objective power: 13 mW. **b**, Signal profiles in the axial plane along the yellow lines in **a. c**, Corrective wavefront in **a. d,g**, Maximum intensity projections of dendrites in the Thy1-YFP-H mouse cerebral cortex at 365-375 μm and 490-513 μm below dura, respectively, under 920 nm excitation, without and with AO (phase modulation). Post-objective power: 31 and 128 mW, respectively. **e,h**, Signal profiles along the purple and blue lines in **d** and **g**, respectively. **f,i**, Corrective wavefronts in **d** and **g**, respectively. Scale bars, 10 μm in **a** and 5 μm in **d** and **g**. Microscope objective: NA 0.8 16× for a and NA 1.05 25× for **d** and **g**.

**Supplementary Figure 1.**
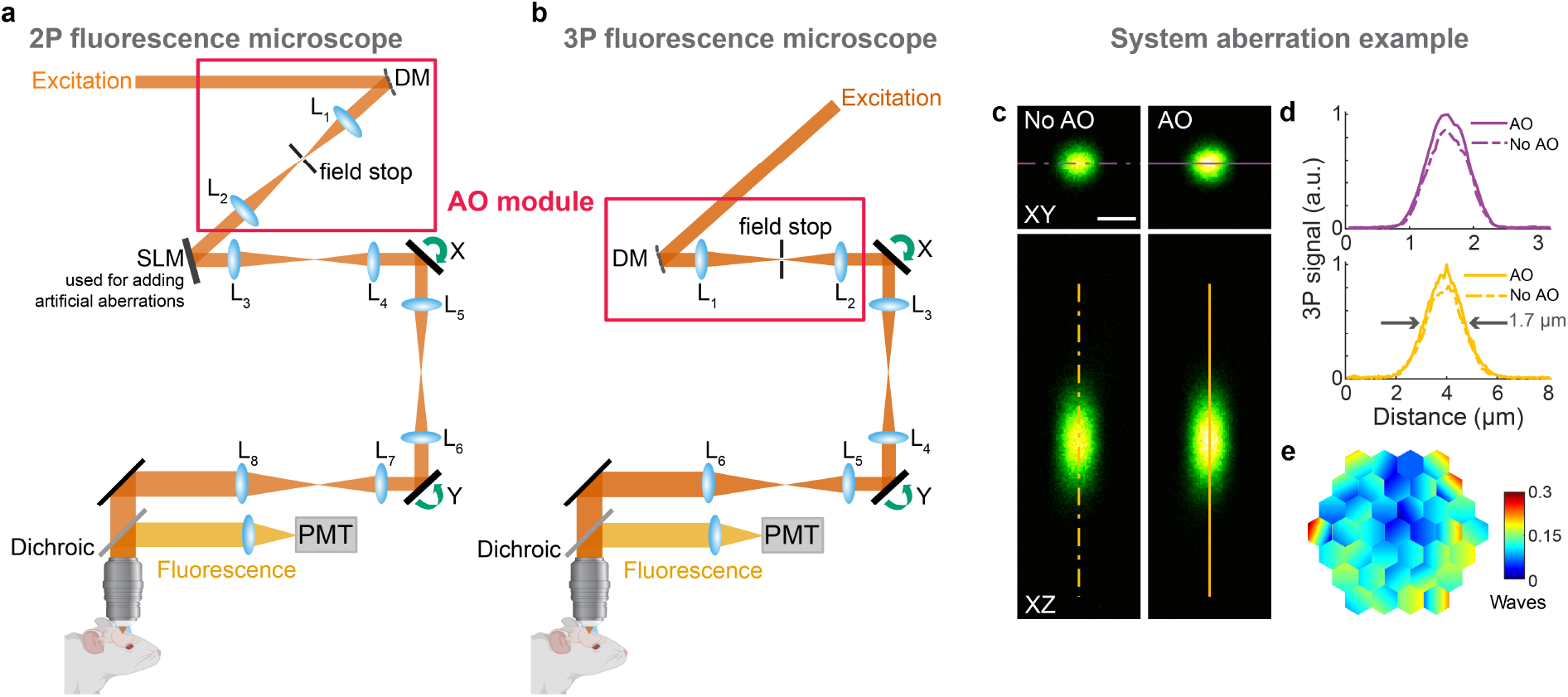
Schematics of AO 2P and 3P fluorescence microscopes, and example system correction. **a, b**, Components of AO 2P and 3P fluorescence microscopes, respectively. DM, deformable mirror; SLM, spatial light modulator (used to introduce artificial aberration); L, lenses; X and Y, galvanometers; PMT, photomultiplier tube. **c**, Lateral and axial 3P images of a 1-μm-diameter red fluorescent bead, under 1300 nm excitation, taken without and with AO. Post-objective power: 0.13 mW. **d**, Signal profiles along the purple and yellow lines in **c. e**, Corrective wavefront applied to the DM. Scale bar, 1 μm. Microscope objective: NA 1.05 25×.

**Supplementary Figure 2.**
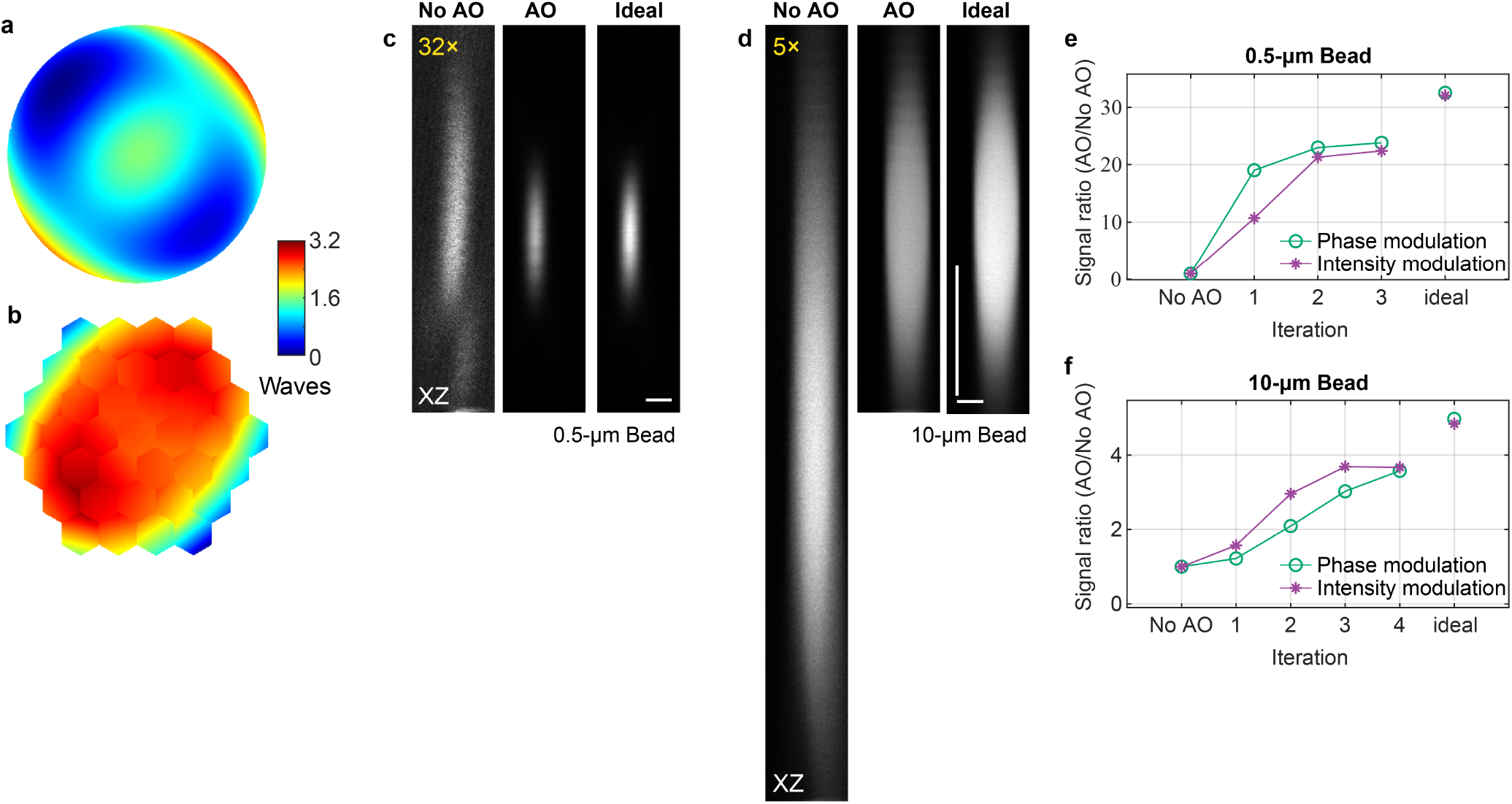
Correcting artificial aberrations for 2P microscopy using phase versus intensity modulation and signal from fluorescent features of different sizes. **a,b**, Artificial aberration introduced with the SLM and corrective wavefront on the DM, respectively. **c,d**, Axial images of 0.5-μm- and 10-μm-diameter fluorescent beads, respectively, without AO, with AO (phase modulation), and under ideal aberration-free conditions (without artificial aberrations applied to the SLM). Digital gains were applied to No AO images to increase visibility. Post-objective power: 6 and 2.3 mW, respectively. **e,f**, 2P signal versus iteration number for 0.5-μm- and 10-μm-diameter beads, respectively, using phase and intensity modulation. Scale bars: 1 μm in **a** and 5 μm in **b**. Microscope objective used: NA 0.8 16×.

Substantial improvements in image quality were also achieved for *in vivo* 2P imaging of the strongly scattering mouse brain through a cranial window. At imaging depths of 370 and 500 μm below dura, aberration correction improved the image signal and contrast of YFP-labeled dendrites (Fig. 1d-i), enabling the identification of fine features such as dendritic spines and axonal boutons – which in some cases can only be resolved in the aberration-corrected image (Fig. 1g,h). Here, and in the examples that follow, the non-corrected (“No AO”) images were taken after correcting aberrations intrinsic to the microscope system (Supplementary Fig. 1c-e) and adjusting the objective correction collar to remove the spherical aberration added by the coverglass used in sample preparation (Methods). These “No AO” images, therefore, represent the best performance that conventional optics and best practice can achieve. The improvements in signal, resolution, and contrast therefore only arose from AO correction of tissue-induced aberrations.

To push the imaging depth even further, we incorporated our AO module into a homebuilt 3P microscope (Supplementary Fig. 1b). After validating its ability to recover nearly-diffraction limited imaging performance under large wavefront aberrations (e.g., beads in a capillary tube, 270× signal improvement, Supplementary Fig. 3), we imaged neuronal structures throughout the mouse cerebral cortex *in vivo*. As an example, at 1300 nm excitation, a single correction pattern applied to the DM drastically improved the image quality of YFP-labeled neurons (Fig. 2a-d, Supplementary Fig. 4, and Supplementary Videos 1 and 2), located 760 μm below dura. Here, AO correction led to 3P signal increases ranging from 7-fold on the cell body to 19-fold on dendrites (Fig. 2c). Moreover, previously invisible synaptic structures (e.g., dendritic spines and spine necks) became easily detectable only after aberration correction (Fig. 2b). It should be noted that, in all mouse brain examples shown here, only 2-3 rounds of correction were used to obtain the final corrective wavefront. Additional rounds did not result in substantial improvement of the fluorescence signal (Supplementary Fig. 5). The data shown is representative of several imaging sessions performed at similar depths (600-870 μm below dura), in different regions of the same animal, as well as in several animals (N=7), and under different labelling conditions (Supplementary Figs. 6–8 and Supplementary Videos 3 and 4). Aberration correction consistently improved the image signal and resolution, allowing us to resolve synaptic structures down to 870 μm below dura (Supplementary Fig. 8), which were otherwise not identifiable without correction.

**Fig. 2.**
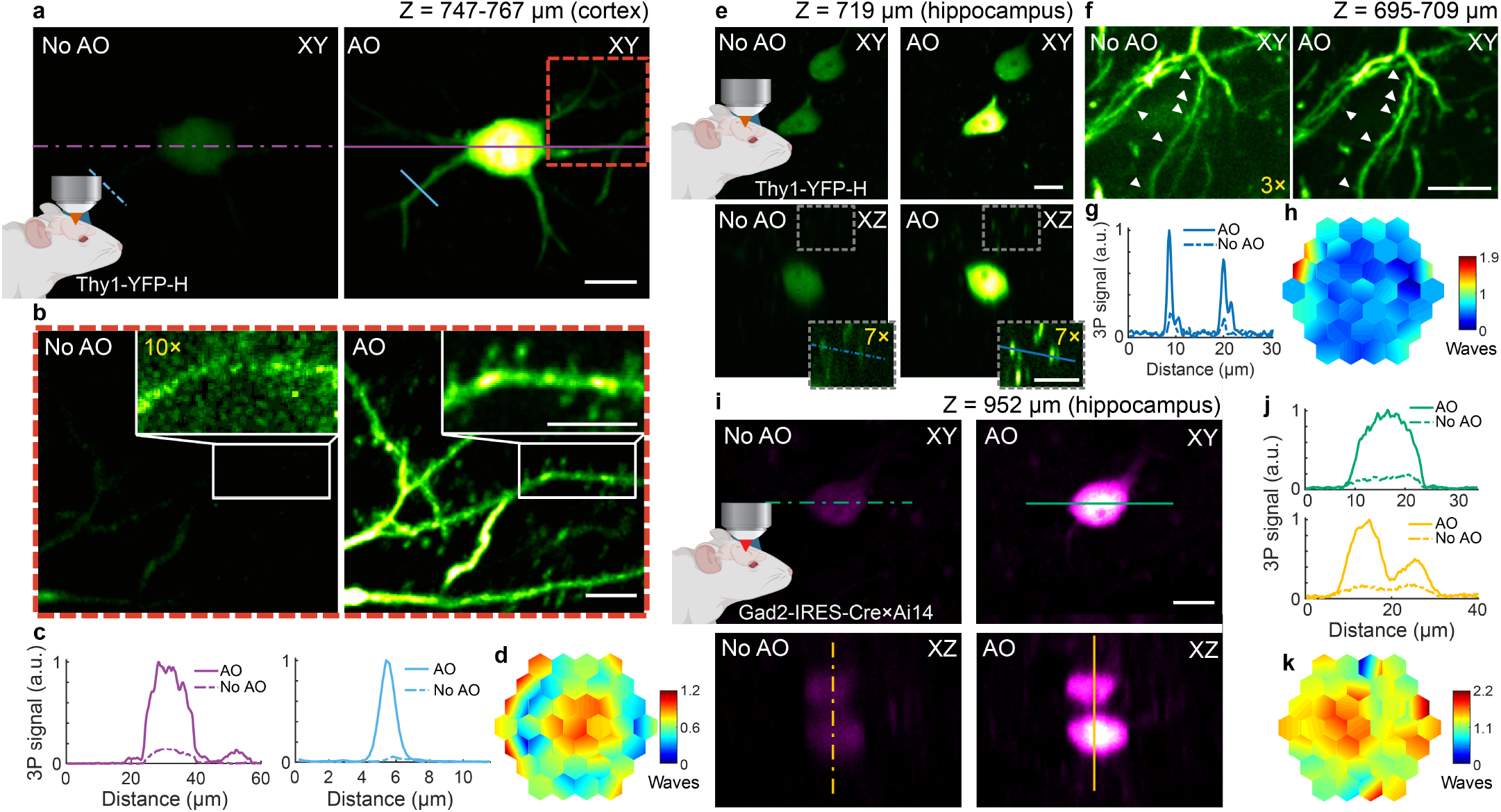
AO enables *in vivo* 3P imaging of cortical and hippocampal neuronal structures in the mouse brain, with subcellular resolution. **a**, Maximum intensity projection (MIP) of a neuron in the mouse cortex (Thy1-YFP-H), at 747-767 μm below dura, under 1300 nm excitation, without and with AO (phase modulation). Post-objective power: 17 mW. **b**, Zoomed-in views of the red square in **a**, at 751-767 μm below dura, without and with AO. Insets, zoomed-in views of the dendrite in white rectangles in **b**. 10× digital gain was applied to No AO inset to increase visibility. Post-objective power: 20 mW. **c**, Signal profiles along the purple and blue lines in **a. d**, Corrective wavefront in **a** and **b. e**, Lateral and axial images of neurons in the mouse hippocampus (Thy1-YFP-H), 719 μm below dura, under 1300 nm excitation, without and with AO (phase modulation). Post-objective power: 16 mW. Insets, zoomed-in views of the gray square in **e**. 7× digital gain was applied to increase visibility. **f**, MIP of neuronal processes above the cell body in **e**, at 695-709 μm below dura, without and with AO. White arrows: dendritic spines. Post-objective power: 26 mW. 3× digital gain was applied to image without AO to improve visibility. **g**, Signal profiles along the blue lines in **e. h**, Corrective wavefront in **e** and **f**. **i**, Lateral and axial images of neurons in the mouse hippocampus (Gad2-IRES-Cre × Ai14 (Rosa26-CAG-LSL-tdTomato)), 952 μm below dura, under 1700 nm excitation, without and with AO (phase modulation). Post-objective power: 30 mW. **j**, Signal profiles along the green and yellow lines in **i. k**, Corrective wavefront in **i**. Scale bars, 10 μm. Microscope objective: NA 1.05 25×.

**Supplementary Figure 3.**
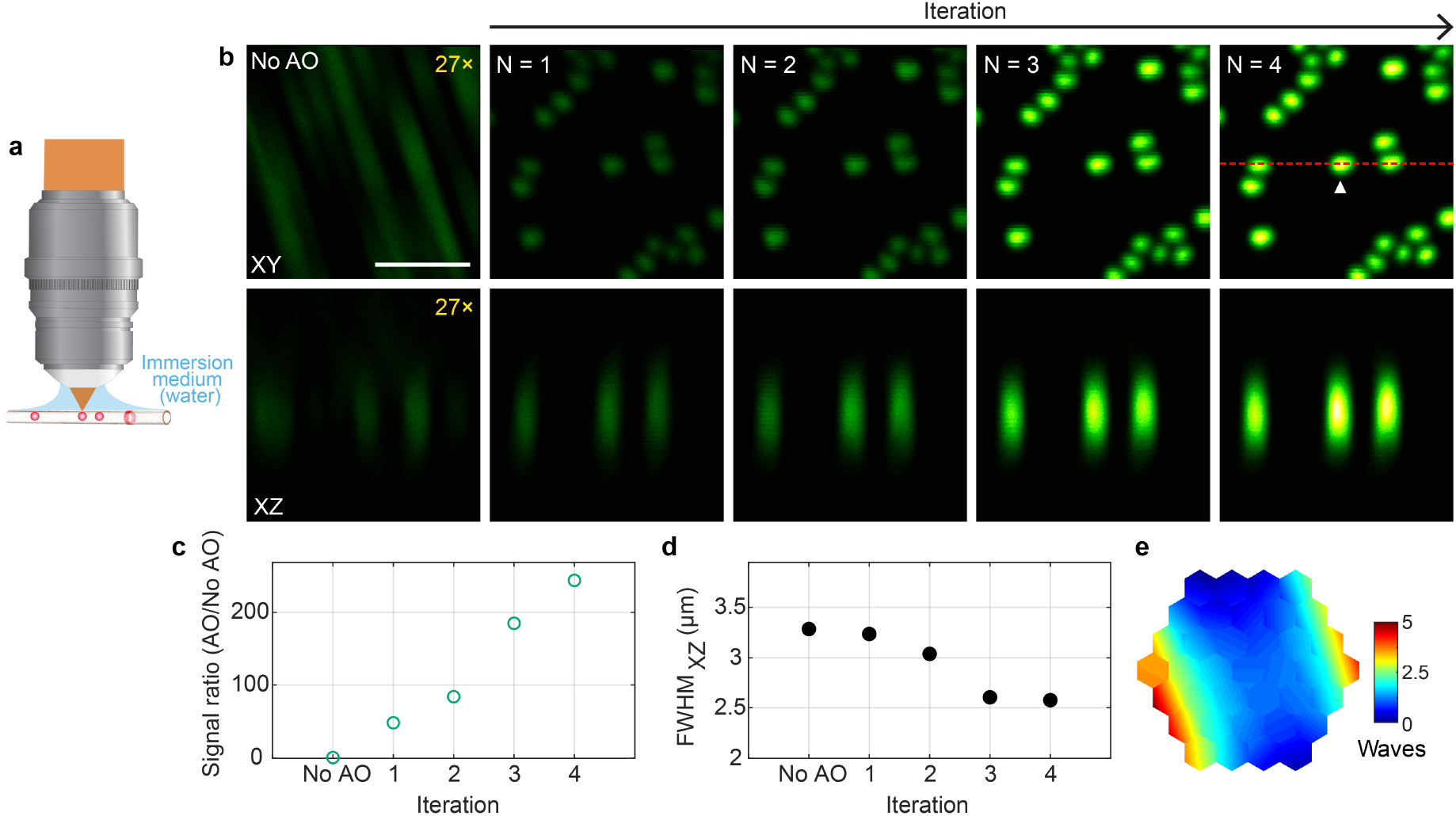
AO improves 3P imaging of beads in a capillary tube. **a**, Schematics of sample geometry of 1-μm-diameter fluorescent beads in an air-filled capillary tube. **b**, Lateral and axial (along red dotted line) images of beads without and with AO (phase modulation). Post-objective power: 0.13 mW. Digital gains were applied to No AO images to increase visibility. **c,d**, 3P signal improvement (AO/No AO) and axial full width at half maximum (FWHM) of a representative bead (white arrowhead in **b**) as a function of the iteration #, respectively. **e**, Corrective wavefront. Scale bar, 5 μm. Microscope objective: NA 1.05 25×.

**Supplementary Figure 4.**
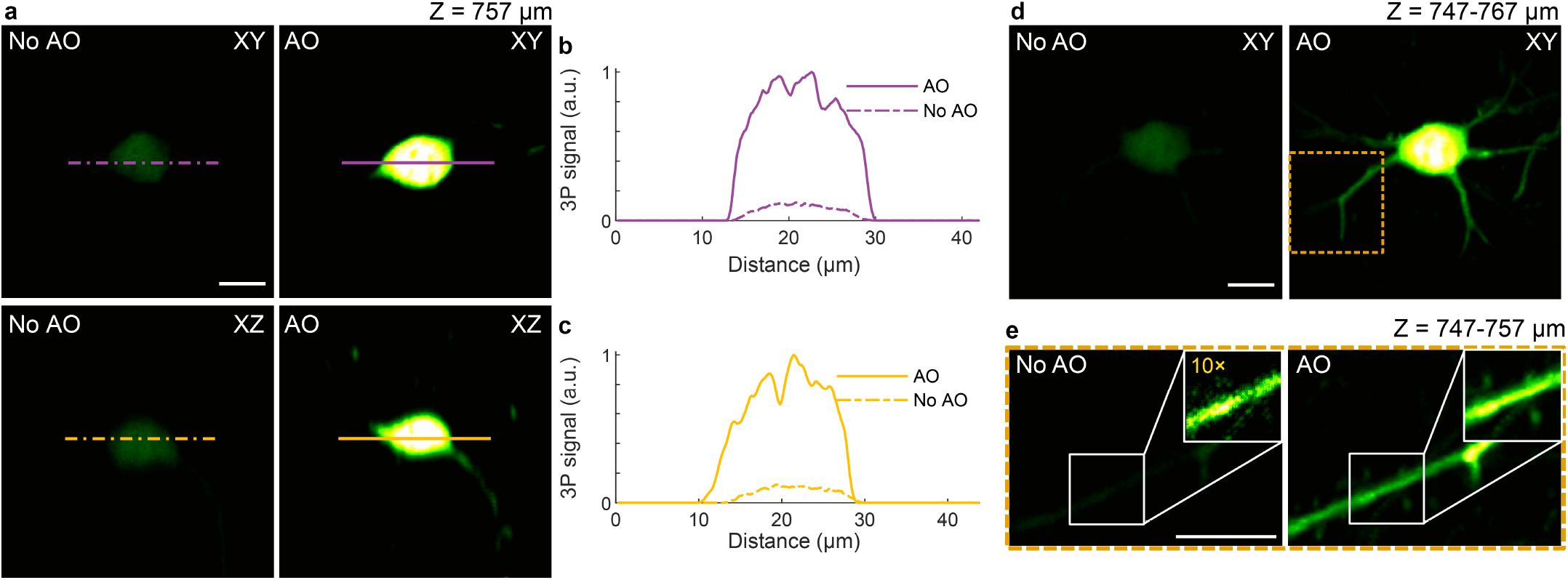
AO improves *in vivo* 3P imaging of cortical neurons in the mouse brain (same cell body as in Fig. 2a). **a**, Lateral and axial images of a neuronal cell body (Thy1-YFP-H), at 757 μm below dura, under 1300 nm excitation, without and with AO. Post-objective power: 17 mW. **b,c**, Signal profiles along the purple and yellow lines in **a. d**, Maximum intensity projection (MIP) of same neuron as in **a**, 747-767 μm below dura, under 1300 nm excitation, without and with AO. Post-objective power: 17 mW. **e**, MIP of the yellow square in **d**, at 747-757 μm below dura, without and with AO. Insets, zoomed-in views of dendrite in white box. 10× digital gain was applied to the inset without AO to improve visibility. Post-objective power: 20 mW. Scale bar, 10 μm. Microscope objective: NA 1.05 25×.

**Supplementary Figure 5.**
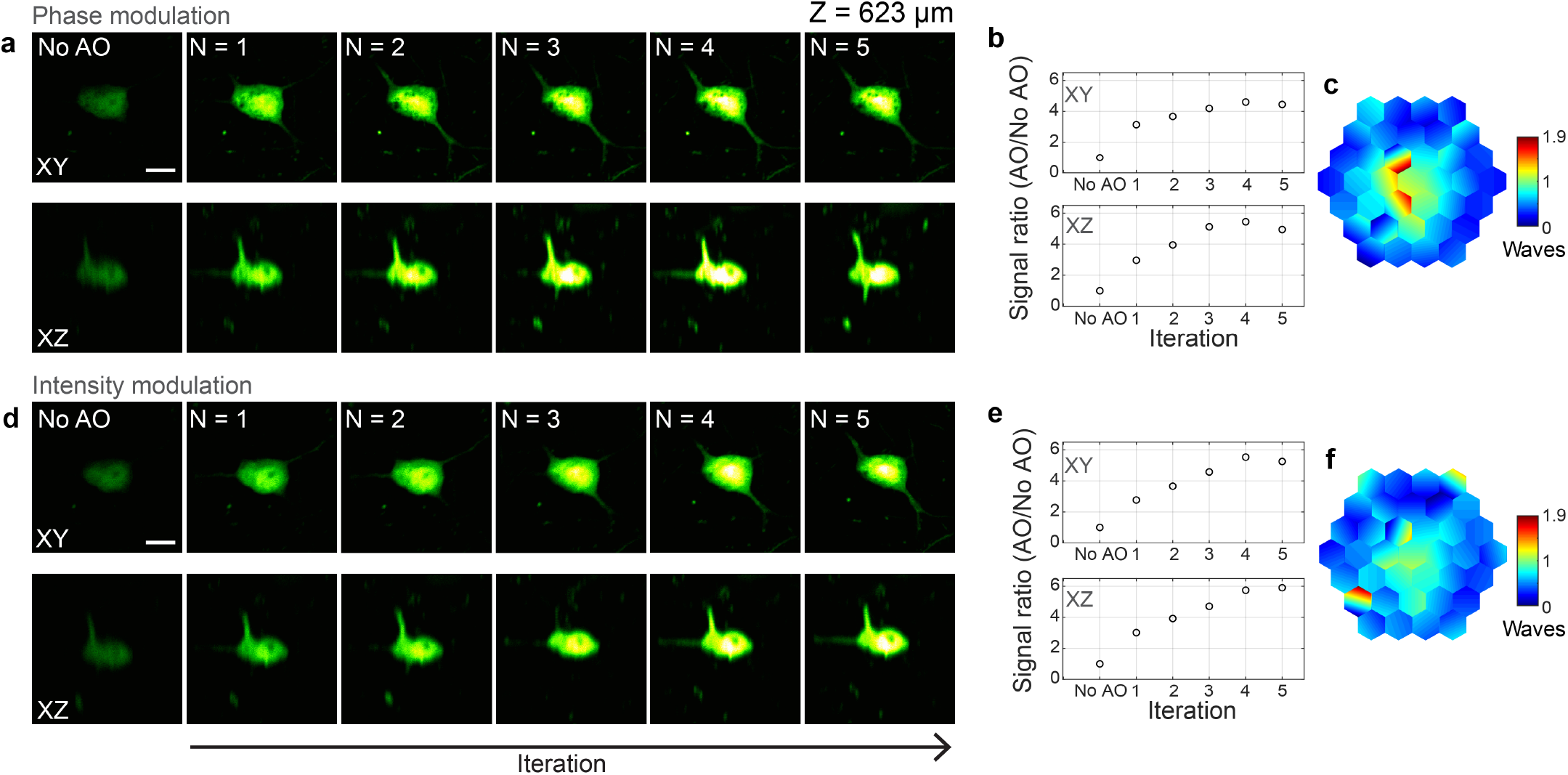
Effect of iterations on 3P fluorescence signal improvement for phase and intensity modulation-based aberration correction in the mouse brain *in vivo*. **a,d**, Lateral and axial images of a neuron in the mouse cortex (Thy1-YFP-H), 623 μm below dura, under 1300 nm excitation, without AO correction and after running aberration measurement a total of N = 1-5 iterations, using phase and intensity modulation, respectively. Post-objective power: 20.8 and 23.6 mW, in **a** and **d** respectively. **b, e**, 3P signal improvement (AO/No AO) with iterations, for phase and intensity modulation, respectively. The plotted signal is the average pixel intensity within a 16×16-pixel area around the image maximum. **c,f**, Corrective wavefronts measured with phase and amplitude modulation, respectively. Scale bars, 10 μm. Microscope objective: NA 1.05 25×.

**Supplementary Figure 6.**
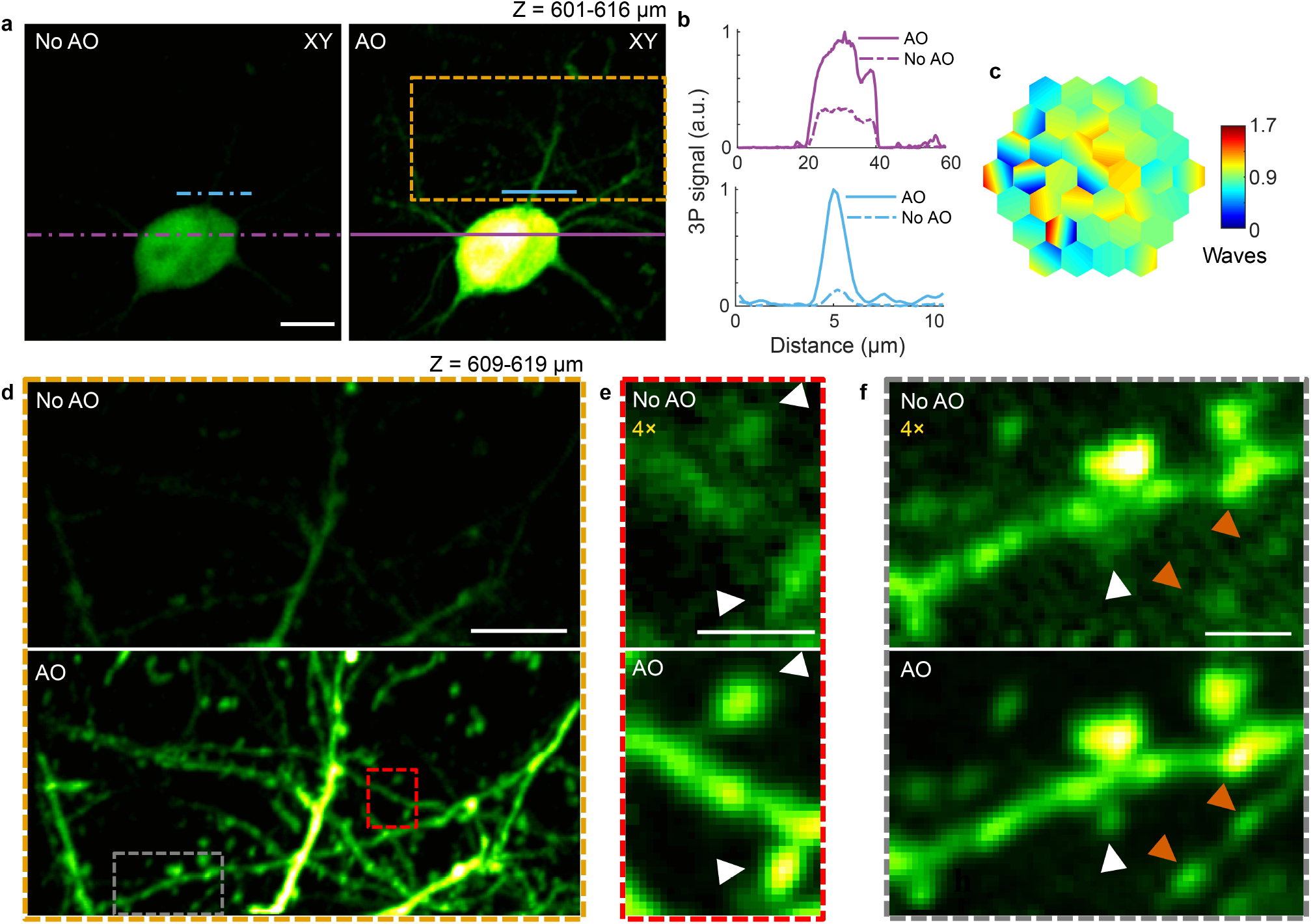
AO enables *in vivo* 3P imaging of dendritic spines and axonal boutons in deep layers of the mouse cortex. **a**, Maximum intensity projection (MIP) of a neuron in the mouse cortex (Thy1-YFP-H), at 601-616 μm below dura, under 1300 nm excitation, without and with AO. Post-objective power: 17 mW. **b**, Signal profiles along the purple and blue lines in **a**. **c**, Corrective wavefront in **a. d**, MIP of the orange box in **a**, at 609-619 μm below dura, without and with AO. Post-objective power: 25.6 mW. **e,f**, Zoomed-in views of the red and gray boxes in **d**, respectively. 4× digital gain was applied to images without AO to improve visibility. White arrowheads: dendritic spines; orange arrowheads: axonal boutons. Scale bars: 10 μm in **a** and **d**; 2 μm in **e** and **f**. Microscope objective: NA 1.05 25×.

**Supplementary Figure 7.**
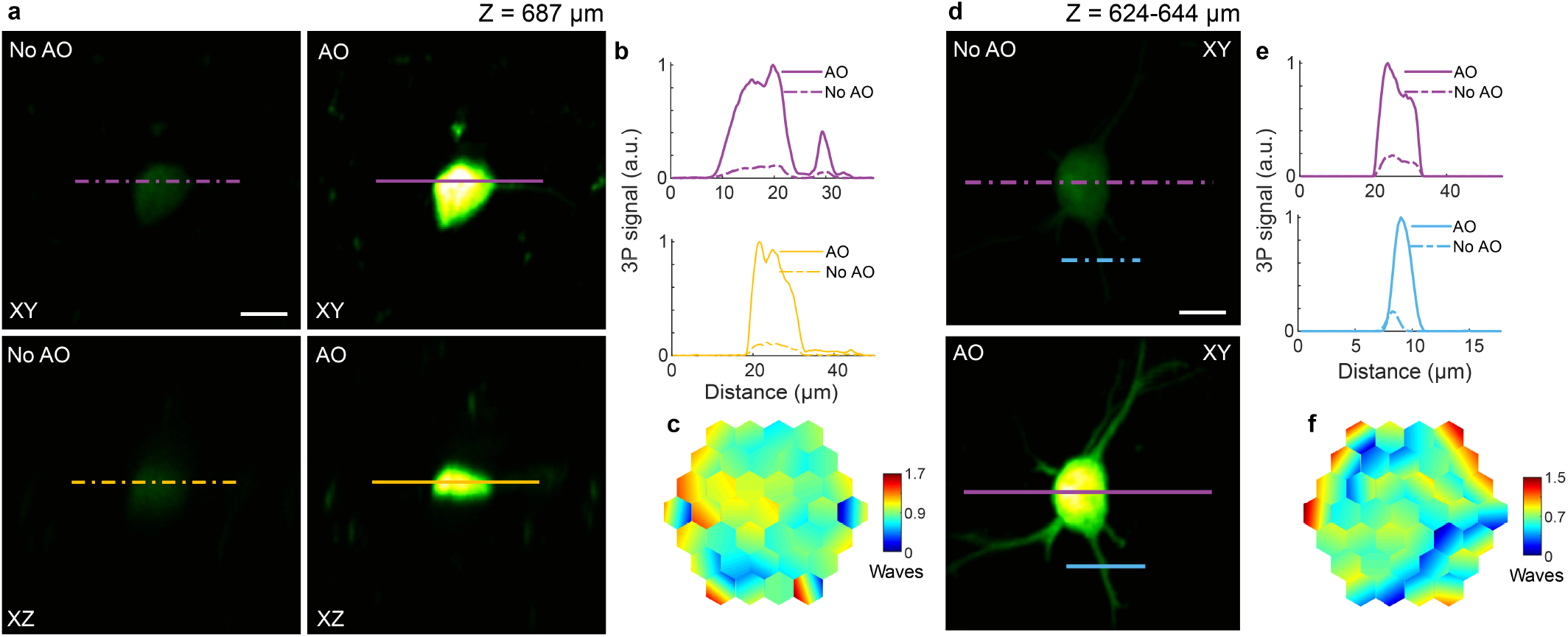
AO improves *in vivo* 3P imaging of cortical neuronal structures in the brain of a Thy1-GFP-M mouse. **a**, Lateral and axial images of a neuron in the mouse cortex (Thy1-GFP-M), at 687 μm below dura, under 1300 nm excitation, taken without and with AO. Post-objective power: 35 mW. **b**, Signal profiles along the purple and yellow lines in **a. c**, Corrective wavefront in **a. d**, Maximum intensity projection of a neuron in the mouse cortex (Thy1-GFP line M, different animal than in **a**), at 624-644 μm below dura, under 1300 nm excitation, without and with AO. Post-objective power: 13 mW. **e**, Signal profiles along the purple and blue lines in **d**. **f**, Corrective wavefront in **d**. Scale bars, 10 μm. Microscope objective: NA 1.05 25×.

**Supplementary Figure 8.**
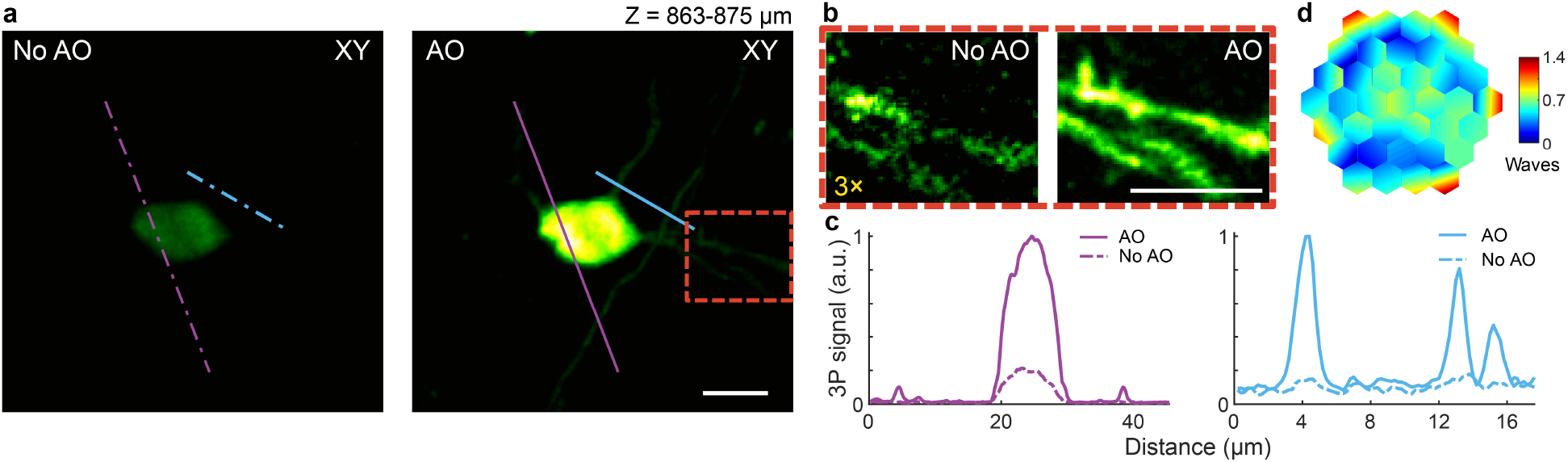
AO enables *in vivo* 3P imaging of synapses in the mouse cortex, 869 μm below dura. **a**, Maximum intensity projection of a neuron in the mouse cortex (Thy1-YFP-H), at 863-875 μm below dura, under 1300 nm excitation, taken without and with AO. Post-objective power: 42 mW. **b**, Zoomed-in views of the red box in **a**. 3× digital gain was applied to the image taken without AO to improve visibility. **c**, Signal profiles along the purple and blue lines in **a**. **d**, Corrective wavefront. Scale bar, 10 μm. Microscope objective: NA 1.05 25×.

Imaging beyond the mouse neocortex and into the hippocampus is difficult owing to the highly scattering white matter. Our AO module enabled imaging of hippocampal structures with subcellular resolution, without the need to perform any invasive procedure (e.g., cortical tissue removal, optical device insertion) other than the cranial window implantation (Supplementary Videos 5-8). Using an excitation wavelength of 1300 nm, aberration correction significantly improved the 3P fluorescence signal (~4-fold) of YFP-labeled somata (Fig. 2e and Supplementary Video 6), with many synaptic structures (e.g., dendritic spines) clearly resolvable only after AO correction (Fig. 2f and Supplementary Video 7). In addition to 3P excitation of green fluorophores using 1300 nm, excitation of red fluorophores with 1700 nm has been shown to improve tissue penetration^1^. To assess the performance of our module under 1700 nm excitation, we imaged tdTomato-labeled neuronal cell bodies at 952-1020 μm below dura, and found a similar improvement in the image signal and contrast after AO correction (Fig. 2i-k, Supplementary Fig. 9, and Supplementary Video 8).

**Supplementary Figure 9.**
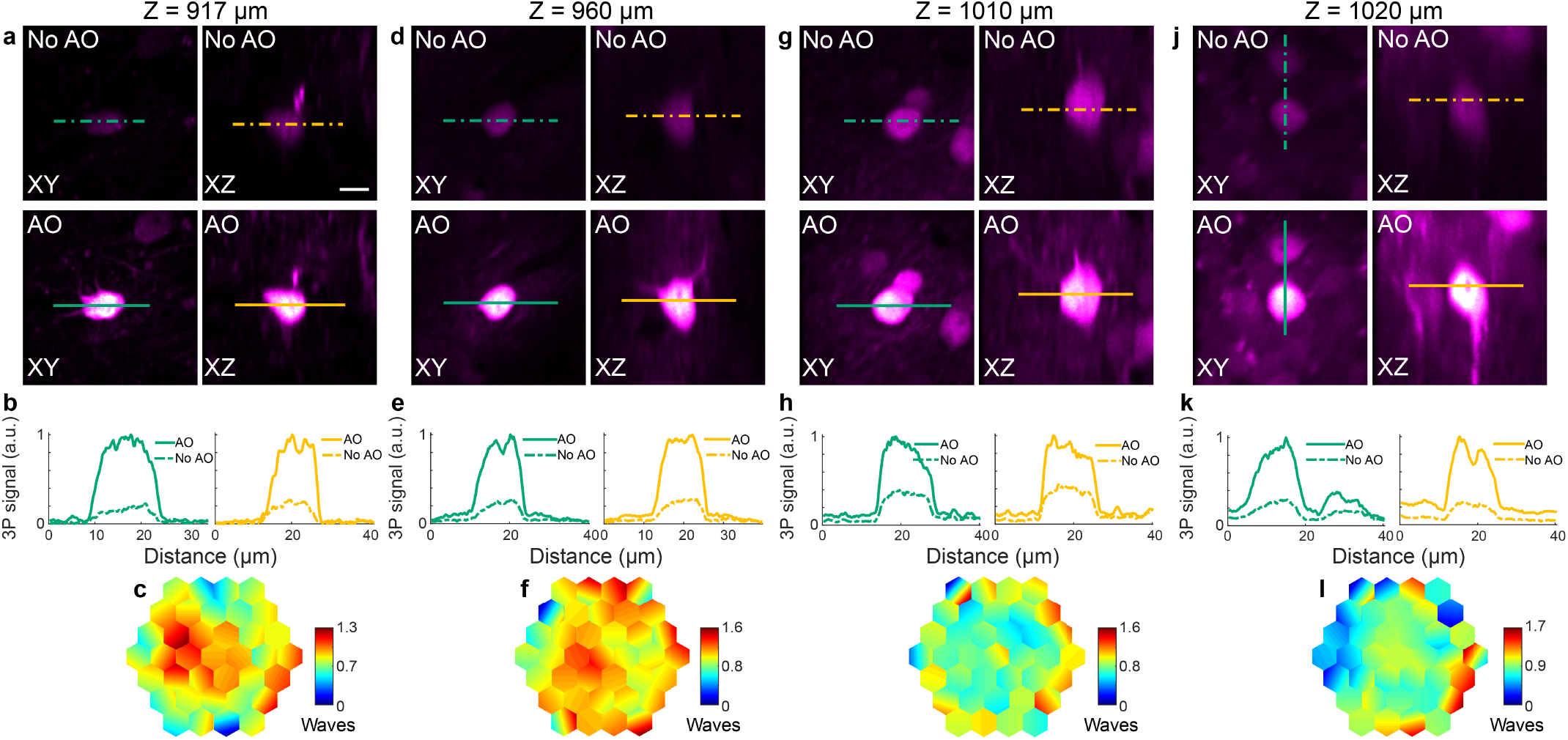
AO improves *in vivo* 3P imaging of hippocampal structures at different depths in the mouse brain, with 1700 nm excitation. **a,d,g,j**, Lateral and axial images of neurons in the mouse hippocampus, without and with AO, at 917, 960, 1010, and 1020 μm below dura, respectively. Post-objective powers: 26.5, 10, 27, and 24 mW, respectively. **b,e,h,k**, Signal profiles along the green and yellow lines in **a, d, g**, and **j**, respectively. **c,f,i,l**, Corrective wavefront in **a, d, g**, and **j**, respectively. For **a**, a Gad2-IRES-Cre × Ai14 (Rosa26-CAG-LSL-tdTomato) mouse was used; for **d, g**, and **j**, neurons in wildtype mice were infected by a mix of AAV-Syn-Cre and AAV-CAG-FLEX-tdTomato. Scale bar, 10 μm. Microscope objective: NA 1.05 25×.

The excellent imaging performance attained by incorporating our AO module into a 3P microscope motivated us to image neuronal structures in an even more challenging environment of the central nervous system: the spinal cord (Fig. 3a-g). The high neuronal density^12^, along with the surface curvature and strong scattering caused by superficial axon tracts, cause wavefront distortions which have prevented 2P microscopy from imaging neuronal structures beyond the most superficial < 200 μm of the spinal cord dorsal horn^13^ – spanning a depth of roughly 500 μm. We performed *in vivo* 3P imaging of GFP-labeled neuronal structures at depths exceeding 400 μm below dura through a dorsal laminectomy^12^, in 9-to 10-week old adult mice, under 1300 nm excitation (Fig. 3a-g). After aberration correction, we achieved signal improvements ranging from 2-to 5-fold (Fig. 3c,f) and more clearly visualized fine neuronal structures.

**Fig. 3.**
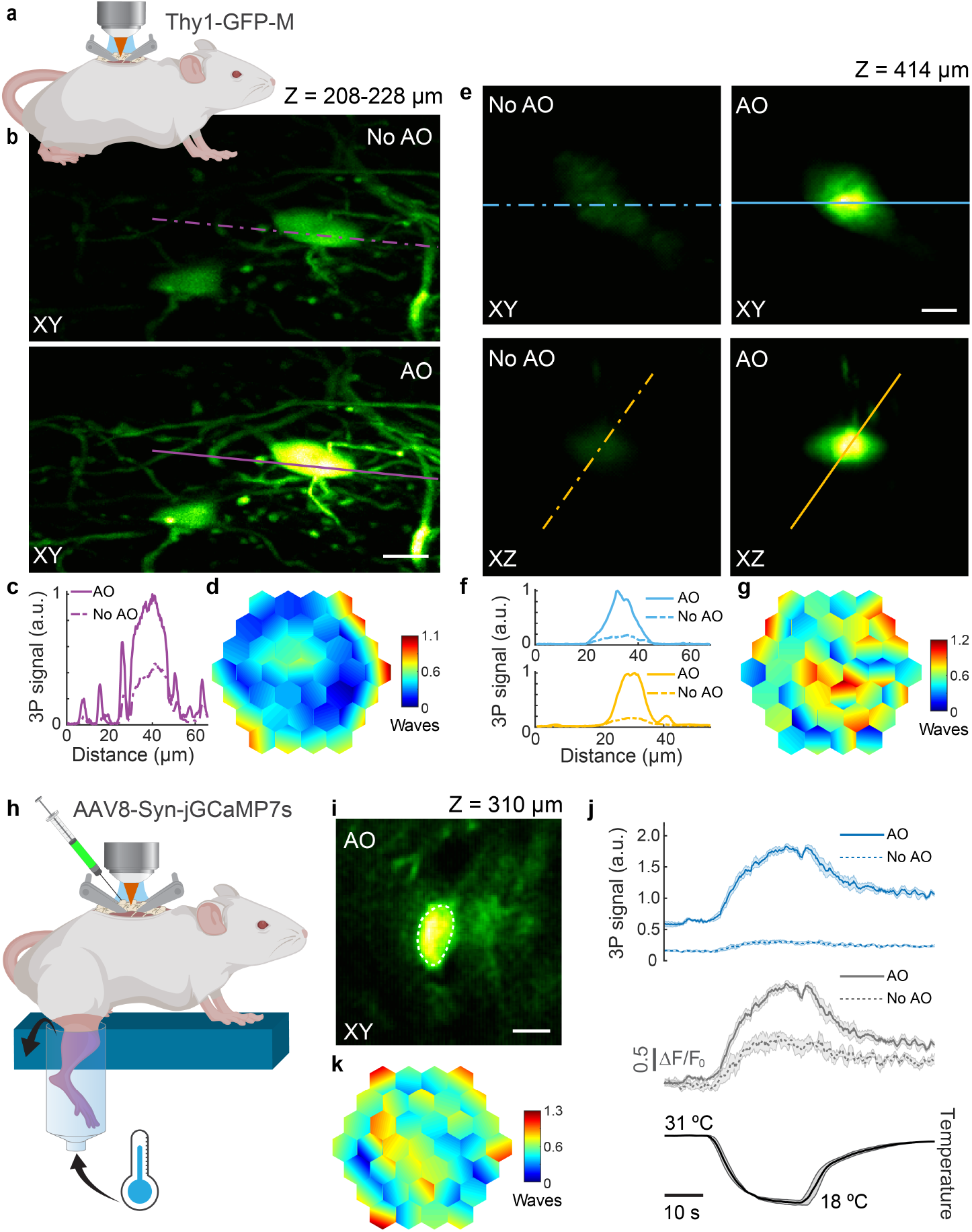
AO improves *in vivo* 3P structural and functional imaging in the mouse spinal cord. **a**, Schematic of *in vivo* imaging in the dorsal horn of the mouse spinal cord. **b**, Maximum intensity projection of spinal cord neurons (Thy1-GFP-M), 208-228 μm below dura, under 1300 nm excitation, without and with AO (phase modulation). Post-objective power: 18.3 mW. **c**, Signal profiles along the purple lines in **b. d**, Corrective wavefront in **b. e**, Lateral and axial images of a neuron (Thy1-GFP-M), 414 μm below dura, under 1300 nm excitation, without and with AO (phase modulation). Post-objective power: 89 mW. **f**, Signal profiles along the blue and yellow lines in **e. g**, Corrective wavefront in **e. h**, Schematic for recording calcium activity in jGCaMP7s-expressing neurons of the dorsal horn in the mouse spinal cord (AAV8-Syn-jGCaMP7s), in response to cooling stimuli applied to the skin of the hindlimb. **i**, Lateral image of a neuron, 310 μm below dura, under 1300 nm excitation, after AO correction. **j**, (top) 3P fluorescence signal and (middle) calcium transients (*ΔF/F_0_*), during (bottom) temperature stimulation, without and with AO (phase modulation), for the neuronal cell body shown in **i**. 4-trial average; shaded area: s.e.m. Post-objective power: 4.2 mW. **k**, Corrective wavefront in **i**. Scale bars, 10 μm. Microscope objective: NA 1.05 25×.

Finally, our AO module enabled us to reliably record somatosensory-evoked calcium transients in the spinal cord dorsal horn at depths beyond 300 μm. Due to its optically challenging nature, such recordings had been limited to the most superficial (< 100 μm) layers, preventing a comprehensive understanding of the complex coding of somatosensory stimuli in the spinal cord circuitry. We recorded calcium transients in jGCaMP7s^14^-expressing spinal cord neurons, in response to cooling applied to the hindlimb skin, at imaging depths where temperature responses had not been previously studied (Fig. 3h-j and Supplementary Video 9)^12^. At an imaging depth of 310 μm below dura, aberration correction substantially improved the signal of a neuronal cell body (by 5.9-fold) and increased the calcium transient amplitudes, with a 2.1-fold increase in the peak calcium-dependent fluorescence change (*ΔF/F*_0_) (Fig. 3j). Although imaging depths exceeding 400 μm were possible, we did not find cooling-responding neurons at such depths.

To summarize, by combining a novel AO module into a 2P and 3P fluorescence microscope, we achieved drastic improvements in image quality, along with subcellular resolution, on a variety of biological structures *in vivo* at great depths. Owing to the higher order nonlinearity of 3P excitation, we observed a particularly large increase in 3P image signal and contrast from correcting tissue-induced aberrations. In the mouse brain, this combination allowed us to resolve synaptic structures in deep cortical layers as well as subcortical areas, otherwise invisible without AO correction, while keeping post-objective average laser powers between ≈10-40 mW (see Supplementary Table 1 for all imaging parameters). In the mouse spinal cord, our AO module enabled subcellular-resolution imaging of neuronal structures, at depths of more than twice of what previous studies had shown^15^. Moreover, aberration correction drastically improved the quality of somatosensory-evoked calcium transient recordings, at depths much beyond (> 3×) what had been previously reported^12,16,17^, opening the door for unravelling previously inaccessible spinal cord circuitry. Besides the ability to achieve subcellular resolution at depth, by improving the focus quality through aberration correction, the excitation power can be significantly reduced, minimizing the out-of-focus background (thus extending the imaging depth limit) and decreasing the power delivered to the sample well below the levels where heating-related effects^18^, or photo-damage^19^, would be a concern.

The high power throughput, ease of implementation, and small footprint needed by our AO module, along with its polarization- and wavelength-independent operation, provides easy integration into existing laser-scanning multiphoton microscopes – such as 2P/3P microscopes, as well as other point-scanning modalities including those based on harmonic generation and Raman scattering. As such, our module can be adopted by a variety of biological laboratories, enabling the investigation of biological processes inside living tissues with subcellular resolution, in fields ranging from neurobiology and cancer biology to plant biology.

## Acknowledgements

We thank J. Wu for help with detection system; the Janelia JET team for designing and assembling the dispersion compensation unit; E. Carroll for help with galvo electronics; S. Chen for surgical assistance; R. Natan, K. Borges, and Q. Zhang for helpful discussions. This work was supported by the Howard Hughes Medical Institute (C.R., A.C., Y.L., W.S., D.E.M., and N.J.); the Burroughs Wellcome Fund under the Career Awards at the Scientific Interface (C.R.); Lawrence Berkeley National Laboratory LDRD 20-116 (J.A.R.); and the Firmenich Next Generation Fund, the Terman Fellowship, NIH grants R01DA045664, R01MH116904, and R01HL150566 (X.C.).

## Author contributions

N.J. conceived and supervised the project. D.E.M., A.C., and N.J. developed the AO control program. T.G.B. performed the DM calibration and provided support on its operation. Y.L. and W.S. performed the mouse brain surgery. M.A.M. performed the mouse spinal cord surgery. M.A.M. built spinal cord temperature stimulation device and C.R. developed control program. A.C. built 2P AO setup and collected 2P data. C.R. built 3P AO setup and collected 3P data in brain and beads. J.A.R. designed new 3P system with input from C.R. J.A.R. and C.R. built new 3P system (used for spinal cord experiments). C.R. and M.A.M. designed and performed 3P imaging experiments in spinal cord. C.R. analyzed data and prepared all figures and supplementary material. X.K.C. supervised spinal cord experiments. C.R. and N.J. wrote the manuscript with feedback from M.A.M. and input from all other authors.

---

**Supplementary Table 1:** Experimental parameters used for the aberration measurement data shown in this work

| Fig. | Imaging depth (μm) | Post-objective power (mW) | Excitation wavelength (nm) | AO iterations | Modulation strategy | mouse line/age/acute or chronic imaging |
| --- | --- | --- | --- | --- | --- | --- |
| 1 a | 110 | 13 | 920 | 4 | phase | zebrafish line: Tg(β-actin:HRAS-EGFP) |
| 1 d | 365-375 | 31 | 920 | 3 | phase | Thy1-YFP-H/>30wks/acute |
| 1 g | 490-513 | 128 | 920 | 3 | phase | Thy1-YFP-H/>30wks/acute |
| 2 a | 747-767 | 17 | 1300 | 3 | phase | Thy1-YFP-H/6wks/acute |
| 2 b | 751-767 | 20 | 1300 | 3 | phase | Thy1-YFP-H/6wks/acute |
| 2 e | 719 | 16 | 1300 | 3 | phase | Thy1-YFP-H/>30wks/chronic |
| 2 f | 695-709 | 26 | 1300 | 3 | phase | Thy1-YFP-H/>30wks/chronic |
| 2 i | 952 | 30 | 1700 | 3 | phase | Gad2-IRES-Cre Jax X Ai14/>30wks/chronic |
| 3 b | 208-228 | 18.3 | 1300 | 2 | phase | Thy1-GFP-M/10wks/acute |
| 3 e | 414 | 89 | 1300 | 3 | phase | Thy1-GFP-M/9wks/acute |
| 3 i | 310 | 4.2 | 1300 | 2 | phase | AAV8-Syn-jGCaMP7s/5wks/acute |
| Supp. Fig. | Imaging depth (μm) | Post-objective power (mW) | Excitation wavelength (nm) | AO rounds | Modulation strategy | mouse line/age/acute or chronic imaging |
| 1 c | N/A | 0.13 | 1300 | 1 | phase | N/A |
| 2 c | N/A | 6 | 920 | 3 | phase | N/A |
| 2 d | N/A | 2.3 | 920 | 4 | phase | N/A |
| 3 b | N/A | 0.13 | 1300 | 4 | phase | N/A |
| 4 a | 757 | 17 | 1300 | 3 | phase | Thy1-YFP-H/6wks/acute |
| 4 e | 747-757 | 20 | 1300 | 3 | phase | Thy1-YFP-H/6wks/acute |
| 5 a | 623 | 20.8 | 1300 | 5 | phase | Thy1-YFP-H/>30wks/chronic |
| 5 d | 623 | 23.6 | 1300 | 5 | intensity | Thy1-YFP-H/>30wks/chronic |
| 6 a | 601-616 | 17 | 1300 | 4 | phase | Thy1-YFP-H/>30wks/chronic |
| 6 d | 609-619 | 25.6 | 1300 | 4 | phase | Thy1-YFP-H/>30wks/chronic |
| 7 a | 687 | 35 | 1300 | 2 | phase | Thy1-GFP-M/>30wks/acute |
| 7 d | 624-644 | 13 | 1300 | 2 | phase | Thy1-GFP-M/5wks/acute |
| 8 a | 863-875 | 42 | 1300 | 2 | phase | Thy1-YFP-H/5wks/acute |
| 9 a | 917 | 26.5 | 1700 | 3 | phase | Gad2-IRES-Cre × Ai14/>30wks/chronic |
| 9 d | 960 | 10 | 1700 | 3 | phase | AAV-Syn-Cre + AAV-CAG-FLEX-ttdTomato/11wks/acute |
| 9 g | 1010 | 27 | 1700 | 3 | phase | AAV-Syn-Cre + AAV-CAG-FLEX-ttdTomato/11wks/acute |
| 9 j | 1020 | 24 | 1700 | 3 | phase | AAV-Syn-Cre + AAV-CAG-FLEX-ttdTomato/11wks/acute |
| 10 | N/A |  |  |  |  |  |
| 11 | N/A |  |  |  |  |  |
| 12 | N/A | 2 | 1300 | N/A | N/A | Thy1-YFP-H/6wks/acute |

## Methods

### Animals

All animal experiments were conducted according to the National Institutes of Health guidelines for animal research. Procedures and protocols on mice and zebrafish were approved by the Institutional Animal Care and Use Committee at Janelia Research Campus, Howard Hughes Medical Institute; and the Animal Care and Use Committee at the University of California, Berkeley. Details on animal preparations are available below.

### Excitation Source

The 2P excitation source was a mode-locked titanium:sapphire laser (Chameleon Ultra II; Coherent) operating at 920 nm. The 3P excitation source consisted of a two-stage optical parametric amplifier (Opera-F; Coherent) pumped by a 40 W diode-pumped femtosecond laser (Monaco 1035-40-40; Coherent) operating at 1035 nm and 1 MHz, providing a broad tuning range (650-920 nm and 1200-2500 nm). Opera-F was operated at 1300 and 1700 nm for 3P excitation, for which the average output power was ~1.5 and ~0.9 W (1.5 and 0.9 μJ per pulse at 1 MHz repetition rate), respectively. For 1300 nm excitation, to reduce the group delay dispersion (GDD) at the sample plane, we used a homebuilt single-prism compressor^20^. After compensation, the pulse duration at the focal plane of the objective was measured to be ~54 fs using an autocorrelator (Carpe, APE GmbH). For 1700 nm excitation, since the GDD is anomalous for many of the glasses and crystals used in our microscope, the resulting negative GDD at the sample plane cannot be compensated for using our prism-based compressor. Instead, the high normal dispersion of ZnSe (from a bulk compressor available inside Opera-F) and silicon (from a 3-mm thick window placed at Brewster’s angle^21^) was used to obtain a pulse duration at the sample of ~70 fs after compensation.

### Adaptive optical microscope setup

Simplified diagrams of our homebuilt 2P microscope and 3P microscope are shown in Supplementary Fig. 1.

For the 2P microscope (Supplementary Fig. 1a), a pair of achromat doublets (AC254-200-B and AC254-300-B; Thorlabs) conjugated the segmented deformable mirror (DM) surface to a liquid-crystal spatial light modulator (SLM; Holoeye, PLUTO-NIR) used to introduce artificial aberrations. The SLM plane was then conjugated to a pair of galvanometers (6215H; Cambridge Technology) that were optically conjugated to each other and the back focal plane of a high-numerical aperture (NA) water-dipping objective (Olympus XLPLN25XWMP2, NA 1.05, 25×; or Nikon CFI LWD, NA 0.8, 16×), using three pairs of achromat doublets (AC254-150-B and AC254-60-B, AC508-080-B and AC508-080-B, AC508-75-C and SLB-50-600PIR1; Thorlabs and OptoSigma).

For the 3P microscope, femtosecond pulses at 1300 or 1700 nm were reflected off a segmented deformable mirror (DM). The DM was conjugated to a pair of galvanometers (6215H; Cambridge Technology) that were optically conjugate to each other and the back focal plane of a high-NA water-dipping objective (Olympus XLPLN25XWMP2, NA 1.05, 25×), using three pairs of achromat doublets (Original 3P system: AC254-100-C and 45-804, AC508-080-C and AC508-080-C, AC508-100-C and SLB-50-600PIR2; Thorlabs, Edmund Optics, and OptoSigma. New 3P system: AC254-400-C and AC254-300-C, SL50-3P and SL50-3P, SL50-3P and TTL200MP; Thorlabs).

For both the 2P and 3P microscopes, a field stop (iris diaphragm, Thorlabs) was located at the intermediate image plane between the DM and the X galvo to block unwanted diffraction orders and light reflected off mirror segments at large tilt angles. To translate the focus axially, the objective was mounted on a piezoelectric stage (P-725.4CD PIFOC; Physik Instrumente). The fluorescence signal was collected by the same objective and reflected from a dichroic beam splitter (FF665-Di02-25×36; Semrock), spectrally filtered (2P: FF01-680/SP, Semrock. 3P: FF01-680/SP, Semrock; together with BLP01-442R-25, Semrock, or ET575lp, Chroma, to block the third-harmonic generation signal at 1300 and 1700 nm, respectively), and detected by a photomultiplier tube (H7422-40 or H10770PA-40; Hamamatsu). A Pockels cell was used for controlling the excitation power (2P: M350-80; 3P: M360-40; Conoptics). For the experiments using 1700 nm excitation, we used D_2_O, instead of H_2_O, for the objective immersion media, because of its much lower absorption^21^. Custom-written software was used for image acquisition.

### Deformable mirror (DM) specifications, calibration, and verification procedure

The deformable mirror in this project is a hexagonal tip-tilt-piston DM (Hex-111-X) manufactured by Boston Micromachines Corporation (Supplementary Fig. 10). The device features 37 hexagonal segments, each anchored to three electrostatic actuators by short attachment posts, a 3.8-mm-diameter clear aperture, and a protective window with anti-reflection coating (400 nm to 1100 nm for 2P experiments and 550 nm to 2400 nm for 3P experiments). We developed it for microscopy application from a 331-segment tip-tilt-piston mirror array originally designed for space-based AO^22,23^. The actuators are independently addressable and can achieve a maximum of 3.5 μm of surface-normal stroke for an input voltage of ~200 V. Tip, tilt, and piston control of mirror segments is achieved by applying command voltages to each of the three actuators. The electromechanical system has no measurable hysteresis. The maximal segment piston motion achievable is 3.5 μm, and the maximal tip and tilt angle achievable is ± 5.7 mrad. However, these maxima are not independent. The range of tip and tilt angle achievable depends on the nominal piston value, and vice versa. Coupling forces and torques are generated on the actuators through their mechanical connection to the mirror segment. As a result, the surface normal deflection at a given actuator post depends not only on the voltage applied to that actuator, but also on the state of its neighboring actuators. Nevertheless, there is a one-to-one mapping between the voltages applied to the three actuators and the resulting tip, tilt, and piston orientations of the segment.

**Supplementary Figure 10.**
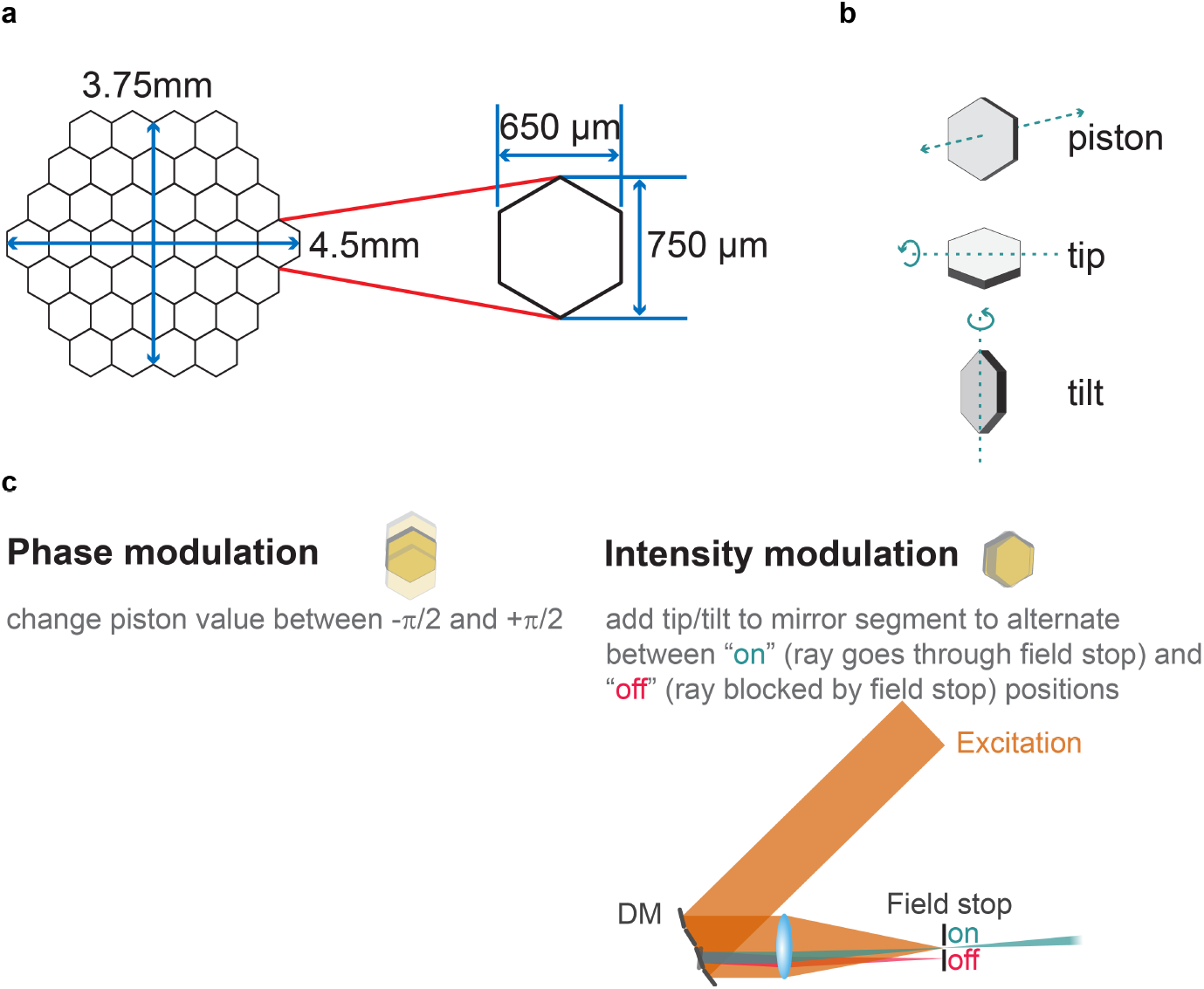
Deformable mirror (DM) specifications. **a**, Diagram of the hexagonal segmented DM (Boston Micromachines Corporation) consisting of 37 segments and a 3.8-mm-diameter clear aperture. **b**, Each segment is controlled by three actuators to provide an angular control of approximately ± 3 mrad in tip and tilt, and a maximum axial stroke (piston displacement) of 3.5 μm. **c**, Diagram showing the two modulation strategies used by our aberration measurement method. See Online Methods for details.

A calibration process was used to fit a mathematical model to that mapping. During calibration, the tip, tilt, and piston of each segment was measured to nanometer-scale precision using a surface mapping interferometer (Zygo 6100) in response to each of 2744 input voltage states applied to the actuators (with voltages ranging from 0 V to 200 V in 14 steps for each actuator). A least squares fourth order polynomial fit was made between the input state (the vector comprised of the three actuator input voltages) and the output state (the vector comprised of the three segment orientations, tip, tilt, and piston). The correlation coefficient of the fit was 1.000. This fit was used to create a lookup table that could be interpolated to map any desired output orientation to a corresponding triplet of required actuator input voltages. The calibration was tested in the interferometer, and yielded precision corresponding to approximately 1 nm in piston motion and 0.1 μradians in tip and tilt.

The calibration of the DM was further tested in our microscope setup. Piston displacement was verified by recording the fluorescence signal from a 1-μm diameter fluorescent bead while piston displacing one mirror segment. A sinusoidal signal trace with constant amplitude and period equal to λ/4 (λ, excitation wavelength) confirmed an accurate calibration and proper functioning of the mirror segment. The same was repeated for all DM segments. Tip/tilt displacement was verified by recording the image of a 10-μm diameter fluorescent bead while scanning a single DM segment along the maximum tip/tilt range. The light reflected off the remaining segments was blocked by the field stop. Using a custom-written MATLAB code, the centroid of the bead was determined for each tip/tilt position, and a plot of measured vs. expected tip/tilt was calculated, taking into consideration the magnification factors in our microscope setup. A slope approximately equal to 1 confirmed accurate calibration of the mirror segment. The same was repeated for all remaining segments. The settling times of the mirror segments were found to be < 100 μs, which allowed us to carry out kHz modulation of the light impinging on it.

### Aberration measurement method

The general aberration measurement procedure was conceptually similar to that described in our previous work^10^. Experimentally, the laser focus was parked at one sample location and the fluorescence signal from this location was used for aberration measurement. The pupil was segmented to 37 regions, corresponding to the number of segments in the DM. The 37 segments were separated into two groups of alternating rows (Supplementary Fig. 11).

First, we fixed the tip, tilt, and piston of one group of 17 segments, but added to each of the other 20 pupil segments a specific tip angle Θ*_i_* and tilt angle *Φ_j_* (*i, j* = 1,2,…*n*) (Supplementary Fig. 11), Step 1), each of which were chosen randomly from an array of *n* angles evenly spaced between-Ψ/2 and Ψ/2. These applied tip/tilt angles caused the beamlet reflecting off the segment to displace along the X and Y axes in the objective focal plane by X*_i_* and *Y_j_* (X*_i_* = *f*tan*(2Θ*_i_/M*) and *Y_j_* = *f*tan*(2Φ*_j_/M*); *f*: focal length of the objective; *M*: magnification from the DM to objective back focal plane), respectively, which changed their interference with the reference focus formed by the other 17 beamlets. With the tip and tilt angles of all segments fixed, using the segments themselves (see details in “Modulation strategies”) we then modulated the phase or intensity of all 20 beamlets, each at a distinct frequency ω_*s*_ (*s* = 1,2,.20), and recorded the fluorescence signal for a time duration T (Supplementary Fig. 11, Step 2). The recorded signal trace was then Fourier transformed (FT) and the Fourier magnitudes at each distinct modulation frequency ω_*s*_ were measured (Supplementary Fig. 11, Step 3), whose values indicated how much individual beamlets interfered with the reference focus at the focal displacement of (X*_i_*, Y*_j_*).

The above procedure was repeated *n* × *n* times, so that all 20 segments sampled the full tip/tilt angles and their corresponding beamlets scanned over a 2D grid in the focal plane with the dimensions 2*f\**tan(Ψ/*M*) by 2*f\**tan(Ψ/*M*) (Supplementary Fig. 11, Step 4). For each beamlet, plotting the Fourier magnitudes versus the displacements (X*_i_*, Y*_j_*), we constructed a 2D map of interference strength of this beamlet with the reference focus at different focal displacements (Supplementary Fig. 11, Step 5). Fitting the map with a 2D Gaussian function, we found the displacements leading to maximal interference between the beamlets and their reference focus, which gave us the tip/tilt angles (i.e., phase gradient) to be applied to this segment in the corrective wavefront.

We then repeated Steps 1-5, but now with the group of 20 pupil segments fixed and forming the reference focus. Modulating the remaining 17 segments, we obtained their 2D maps of interference strength versus displacement (Supplementary Fig. 11, Step 6) and the phase gradients required to shift the corresponding beamlets to coincide with the reference focus. The total fluorescence signal acquisition time was 2 × *n* × *n* × T.

With all the beamlets intersecting at the same location, the next step was to determine the phase offsets that would allow them to constructively interfere^24^. Consider the case of finding the phase offset that enables two beamlets to constructively interfere. With one beamlet assigned as the reference (with unknown phase *θ_r_*), by incrementally adding a phase offset *Δθ* to the other beamlet (with unknown phase θ_1_ at step size ω (i.e., Δθ = ωt), the intensity variation can be described by *I* = 2 + 2cos (ωt+ θ_1_ - *θ_r_*). The phase offset that gives the maximal intensity (i.e., constructive interference) is thus the opposite of the phase of the function cos (ωt + θ_1_ - *θ_r_*). One approach is by Fourier-transforming the time-dependent signal and reading out the phase at frequency ω/2π. To determine the phase offsets of 37 segments, we employed the concept of multidither coherent optical adaptive technique^11^,^25–27^. We modulated the phases of the first 20 rays by piston-displacing each corresponding mirror segment at a distinct frequency *ω_s_* while keeping the phases of the remaining rays, which formed a reference focus, constant. We then Fourier transformed the recorded fluorescence trace (Supplementary Fig. 11, Step 7), and read out the phase offsets that would lead to constructive interference with the reference focus at the modulation frequencies ω*_s_* (*s* = 1,2,…20) (Supplementary Fig. 11, Step 8). We next modulated the phases of the remaining 17 segments while keeping the phases of the first 20 segments unchanged and found the phase offsets for these 17 segments required to ensure constructive interference among beamlets.

With all beamlets intersecting and constructively interfering at the focal plane, we obtained the final corrective wavefront and applied it to the DM for aberration correction. Because the reference foci used for both phase gradient measurements (Supplementary Fig. 11, Step 1-6) and phase offset measurements (Supplementary Fig. 11, Step 7-9) were aberrated to begin with, for larger aberrations, the whole procedure was iterated, as needed, to achieve optimal aberration correction (Supplementary Fig. 11, Step 10).

**Supplementary Figure 11.**
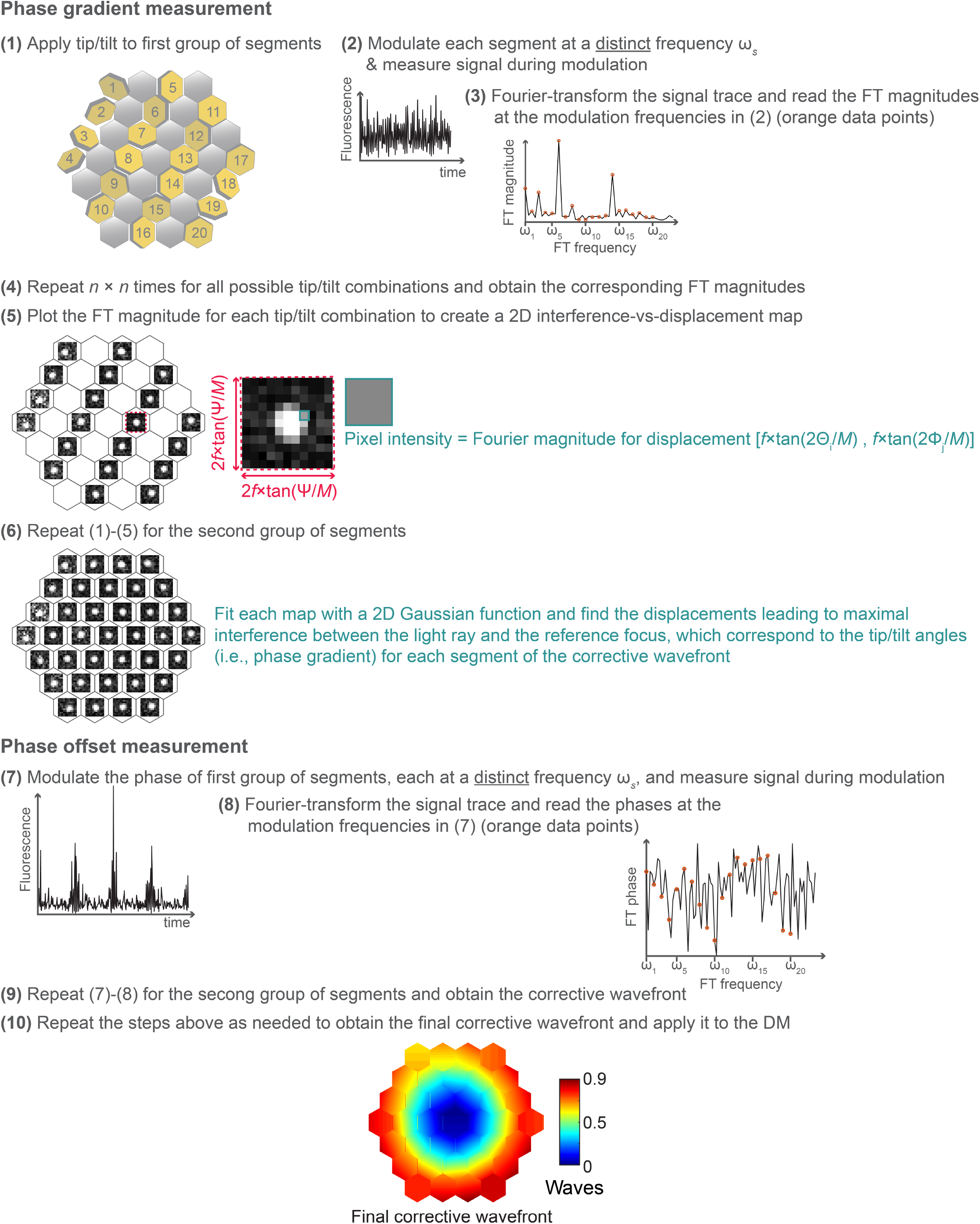
Schematics of the aberration measurement method. See detailed description in Methods.

### Modulation strategies

Two different modulation strategies were used: phase modulation and intensity modulation. During phase modulation, mirror segments were piston-displaced between -π/2 and π/2 (imparting the beamlets reflecting off them a phase shift ranging from -π to π) at frequency ω_*s*_, resulting in a modulation on the detected fluorescence signal. During intensity modulation, a large tip (or tilt) angle was intermittently applied to the mirror segments being modulated, each at a unique ω_*s*_, such that the beamlets impinging on them alternate between two positions: the “on” position, where beamlets went through the field stop placed after the DM; and the “off” position, where beamlets were blocked by the field stop (Supplementary Fig. 10).

During phase modulation, the modulation in the fluorescence signal originates from the varying phase offsets between the modulated rays and the reference focus. In contrast, during intensity modulation, the periodic variation in the fluorescence signal comes from the changes in excitation power at the focal plane. While for phase modulation there is always a maximum present in 2D maps of interference strength versus displacement, for intensity modulation, if the relative phase between the ray and the reference focus is near π/2, a clear maximum would be absent from the resulting map, in which case a π/2 phase offset needs to be added to the corresponding segment and the measurement redone.

Depending on the sample, fluorescent features of different sizes may be used for aberration measurement. We systematically tested and compared the performance of phase versus intensity modulation, each of which was used to measure artificial aberrations using the signal from fluorescent features of different sizes (Supplementary Fig. 2). Experimentally we found that, for 2P microscopy, phase modulation outperformed intensity modulation when signal from small features (< 4 μm) was used for aberration measurement, whereas intensity modulation led to faster improvement of image quality for features larger than 4 μm. For 3P microscopy, phase modulation was found to perform better than or equally with intensity modulation for all feature sizes tested (from 1-μm diameter fluorescent beads to a fluorescent solution). Similar performance between phase and intensity modulation was validated *in vivo* by performing the correction using a neuronal cell body (Supplementary Fig. 7).

### Typical operation parameters and imaging considerations

An example set of operation parameters used for aberration measurement in the mouse brain *in vivo* (Fig. 2) included an integration time of T = 90 ms for each of the 11 × 11 (*n* × *n*) tip/tilt angles, which scanned the modulated rays over 19 μm × 19 μm 2D grids in the focal plane. The overall fluorescence acquisition time was therefore 21.8 s (2 × *n* × *n* × T). Additionally, for the phase measurement portion of the algorithm, the total fluorescence acquisition time was 1.8 s (360 ms per iteration, with a total of 5 iterations). Additional hardware (e.g., DM settling time) and software overheads added to the fluorescence acquisition time and the overall time used for aberration measurement and correction was 3-4× the fluorescence acquisition time. For data in Fig. 2, 3 rounds of aberration measurements were performed to obtain the final corrective wavefront.

“No AO” images were taken after system aberration correction (Supplementary Fig. 1c-e), as well as the adjustment of the objective correction collar to compensate for the spherical aberration introduced by the glass windows overlaying the mouse brain and spinal cord. Furthermore, to minimize additional aberration modes that arose from a tilted window and sample (e.g., coma)^28^, we used the third-harmonic generation signal from the window-tissue interface to determine the angle of the window (Supplementary Fig. 12) and adjusted the mouse using a 2D goniometer stage till the window was perpendicular to the excitation beam. These procedures constitute the best practice and lead to the best performance that conventional optics could achieve. The deterioration in image signal and contrast observed in our “No AO” images, therefore, resulted from tissue-induced aberrations exclusively, and could only be compensated by AO.

**Supplementary Figure 12.**
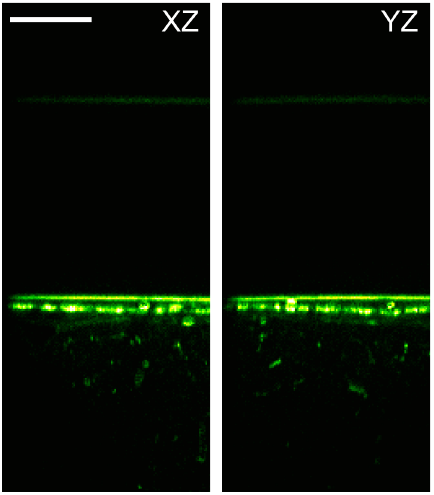
Cranial window alignment using the third-harmonic signal generated at the glass window - brain interface. Axial third-harmonic generation images of a cranial window above a Thy1-YFP-H mouse brain *in vivo.* Both the 3P fluorescence and the third-harmonic signals are shown, where the third-harmonic signal generated at the interfaces of the coverglass are clearly distinguished. Excitation wavelength: 1300 nm (same imaging session as in Fig. 2 a-d). Post-objective power: 2 mW. Scale bar, 50 μm. Microscope objective: NA 1.05 25×.

### Digital image processing

Due to brain motion at depth, the StackReg^29^ image registration plug-in in ImageJ was used for rigid registration in 2D. The “smooth” function from ImageJ that replaced the value of each pixel with the average of 3 × 3 pixels centering on this pixel was applied to all 3P images. 2P images were presented using the “Green hot” lookup table in ImageJ; 3P images were presented with the “Green hot” and “Magenta hot” lookup tables for 1300 and 1700 nm excitation, respectively. For images where the signal was too weak before AO correction, a linear scaling factor was applied to all pixel values to improve visibility with the scaling factor listed on the image. The images presented did not undergo any other digital manipulation.

### Bead samples

Carboxylate-modified fluorescent microspheres (Fluosphere™; Invitrogen) were immobilized on poly(l-lysine)-coated microscope slides (12-550-12, Fisher Scientific).

### Zebrafish preparation

Zebrafish procedures have been described previously^30^. Briefly, Tg(β-actin:HRAS-EGFP) zebrafish embryos (*Danio rerio*) were grown at 28 °C in E3 zebrafish embryo medium. To generate optically transparent embryos, melanin synthesis was inhibited by transferring the embryos into 1× phenylthiourea (PTU) solution in E3 medium, 10-16 hours post fertilization. Before imaging, the chorions were manually removed with forceps under a stereomicroscope. Four-day-old larvae were then anesthetized in E3 medium containing 1× tricaine and immobilized on a dish by embedding them in 0.5% low-melting point agarose with 1× PTU and 1× tricaine. During imaging, E3 medium containing 1× PTU and 1× tricaine was used as immersion medium.

### Mouse preparation (brain imaging)

Cranial window implantation procedures were performed^31^, using aseptic technique, on mice that were anaesthetized with isoflurane (1–2% by volume in O2) and given the analgesic buprenorphine (SC, 0.3 mg per kg of body weight). For 2P imaging experiments, a 3.5-mm diameter craniotomy was made over V1 with dura left intact. A glass window made of two coverslips (Fisher Scientific, thickness no. 1.5) bonded with ultraviolet cured optical adhesives (Norland Optical Adhesives 61) was embedded in the craniotomy and sealed in place with dental acrylic. For 3P imaging experiments, a 5-mm diameter craniotomy was made over V1, with dura left intact. The glass window consisted of a donut-shaped coverslip (inner diameter 4.5 mm, outer diameter 5.5 mm; Potomac Photonics) bonded with ultraviolet cured optical adhesives (Norland Optical Adhesives 61) on top of a 5-mm diameter coverslip (Denville Scientific, thickness no. 1). The window was embedded in the craniotomy and sealed in place with dental acrylic. A titanium head-post was attached to the skull with cyanoacrylate glue and dental acrylic. Acute imaging was performed an hour after the surgery; chronic imaging happened at least one week after the surgery. Mice were head-fixed and anesthetized using isoflurane (1–2% by volume in O2) during imaging.

Some mice also underwent virus injection procedures as described previously^31^. For the examples shown in Supplementary Fig. 9, neurons in the hippocampus were infected with a mixture of AAV-Syn-Cre (10× dilution from 1.8 × 10^13^ GC/mL) and AAV-CAG-FLEX-tdTomato (3.3 × 10^13^ GC/mL) in a wild-type mouse (C57BL/6J). Three injection sites were chosen (AP: −1.7 mm, ML: +1.5 mm; AP: −2.0 mm, ML: +2.0 mm; AP: −2.3 mm, ML: +2.5 mm), at five different depths (0.6, 0.8, 1, 1.2, and 1.4 μm). 100 nL of viral solution were injected at each spot. A cranial window was implanted, as described above, 14 days after virus injection and acute imaging was then performed.

### Mouse preparation (spinal cord imaging)

Acute spinal cord windows were prepared as described previously^12^. Briefly, mice were anesthetized with two successive intraperitoneal injections of 1mg/kg body weight urethane each, 30 mins apart. A tracheotomy was performed to prevent asphyxiation. The T11-13 vertebrae were exposed and stabilized using spinal clamps (STS-A, Narishige). A dorsal laminectomy was performed at T12 to expose the spinal cord. After a wash with Ringer solution (135 mM NaCl, 5.4 mM KCl, 5 mM HEPES, 1.8 mM CaCl_2_, pH 7.2), the spinal cord was covered with a glass window made of a single coverslip (Fisher Scientific No. 1.5). The window and the surrounding custom imaging chamber were stabilized using 2% agarose in Ringer solution. Blood flow through the central blood vessel was continuously monitored throughout the imaging experiment to ensure tissue health. For the functional imaging experiments, wild-type (C57BL/6J) mice had previously been intrathecally injected with ~5 × 10^10^ GC of AAV8-Syn-jGCaMP7s.

### Temperature stimulation (spinal cord imaging)

The left hind limb of the mouse was dehaired and gently fixed inside a custom-designed stimulation device as previously described^12^. The entire limb was continuously exposed to a homogeneous high flow-rate water flow of variable temperature. For each trial, the spinal cord was imaged for a total of 70 s. The first ~10 s were imaged at 31 °C baseline water to obtain the baseline fluorescence and noise. For the following ~30 s the flow was switched to water that was pre-incubated at colder temperatures with the same flow rate. For the remaining ~30 s the flow was switched back to the 31°C baseline water. At least another 120 s passed before the next trial was performed. The actual temperature inside the stimulation chamber was monitored and recorded using a microprobe thermometer (BAT-12, Physitemp) with a Type-K thermocouple placed right next to the mouse’s limb, simultaneously with the fluorescence images. The electric valves controlling the water flow switch were triggered by and synchronized with the image acquisition system.

### Analysis of calcium imaging data

To account for motion in the spinal cord caused by respiration, we processed the image sequences using an iterative cross-correlation-based registration algorithm^31^. We averaged across trials, manually selected the region of interest, and calculated the mean fluorescence within this region. From this, we calculated the fractional change in 3P fluorescence (*ΔF/F_0_*) due to neural activity, with *F*_0_ being the baseline fluorescence calculated as the mean fluorescence during exposure to the baseline temperature (initial ~10 s of the temperature stimulation). For the traces shown in Fig. 3j, we performed a 5-frame moving average.

## Supplementary Videos

The data presented here did not undergo any smoothing or any other digital manipulation, except for image registration.

Supplementary Video 1: ***In vivo* image stacks of cortical neuron in Thy1-YFP-H mouse measured without and with AO**. Same data as shown in Fig. 2a. Scale bar, 10 μm.

Supplementary Video 2: **Zoomed-in views of *in vivo* image stacks of cortical neuron dendrites in Thy1-YFP-H mouse measured without and with AO**. Same data as shown in Fig. 2b. Scale bar, 10 μm.

Supplementary Video 3: ***In vivo* image stacks of cortical neuron in Thy1-YFP-H mouse measured without and with AO**. Same data as shown in Supplementary Fig. 6a. Scale bar, 10 μm.

Supplementary Video 4: **Zoomed-in views of *in vivo* image stacks of cortical neuronal processes with dendritic spines and axonal boutons in Thy1-YFP-H mouse measured without and with AO**. Same data as shown in Supplementary Fig. 6d. Scale bar, 10 μm.

Supplementary Video 5: **Stack of 3P fluorescence and third-harmonic images going from cortex to the hippocampus of a Thy1-YFP-H mouse brain *in vivo*.** This image sequence of combined 3P fluorescence and third-harmonic signals was obtained from 250 μm to 754 μm below the dura of a mouse whose hippocampal neurons are shown in Fig. 2e-h Note the transition from cortex to hippocampus, through the white matter. Scale bar, 20 μm.

Supplementary Video 6: ***In vivo* image stacks of hippocampal neuron in Thy1-YFP-H mouse measured without and with AO**. Same data as shown in Fig. 2e. Scale bar, 10 μm.

Supplementary Video 7: **Zoomed-in views of *in vivo* image stacks of hippocampal neuronal processes in Thy1-YFP-H mouse measured without and with AO**. Same data as shown in Fig. 2f. Scale bar, 10 μm.

Supplementary Video 8: ***In vivo* image stacks of hippocampal neurons in Gad2-IRES-Cre × Ai14 (Rosa26-CAG-LSL-tdTomato) mouse measured without and with AO**. Same data as shown in Fig. 2i. Scale bar, 10 μm.

Supplementary Video 9: **AO improves calcium activity recordings in the mouse spinal cord *in vivo,* 310 μm below dura.** Image sequence corresponding to data shown in Fig. 3i-k (4-trial average). Scale bar, 10 μm.

